# Rational protein engineering to enhance MHC-independent T cell receptors

**DOI:** 10.1101/2024.02.22.581571

**Authors:** Ju-Fang Chang, Jack H. Landmann, Tien-Ching Chang, Yangdon Tenzin, John M. Warrington, Mehmet Emrah Selli, Julie Ritchey, Yu-Sung Hsu, Michael Slade, Deepesh Kumar Gupta, John F. DiPersio, Nathan Singh

**Author notes:** Correspondence: Nathan Singh MD, 660 S. Euclid Avenue, Saint Louis Missouri 63110.

## Abstract

Chimeric antigen receptor (CAR)-based therapies have pioneered synthetic cellular immunity against cancer, however remain limited in their scope and long-term efficacy. Emerging data suggest that dysregulated CAR-driven T cell activation causes T cell dysfunction and therapeutic failure. To re-engage the endogenous T cell response, we designed hybrid MHC-independent T cell receptors (miTCRs) by linking antibody variable domains to TCR constant domains. While functional, we observed stark differences in miTCR-driven T cell function that were dependent on receptor orientation. Using predictive structural modeling, we observed significant biochemical conflicts at the hybrid variable-constant domain interface. To overcome this, we performed iterative sequence modifications and structural modeling to design a panel of miTCR variants predicted to have improved interface stability. Functional screening nominated a variant with superior efficacy to all other miTCRs as well as a standard CAR against high burdens of leukemia.

**Statement of Significance:** Improving the durability of engineered T cell immunotherapies is critical to enhancing efficacy. We used structure-informed design to evolve MHC-independent T cell receptors that drive improved tumor control. This work underscores the central role of synthetic receptor structure on T cell function and provides a framework for improved receptor engineering.

## Introduction

Chimeric antigen receptor (CAR) engineered T cells have paved the path for synthetic cellular immunity against cancer. Durable remission is now realistic for many patients with highly refractory, aggressive B cell cancers using CD19-targeted CAR therapies. Despite these watershed successes, engineered immune cells have yet to meet their potential. Standard-of-care CAR therapies remain restricted to B cell and plasma cell cancers, and far too few patients experience long-term remission. Beyond this, no CAR-based products have demonstrated reliable efficacy against solid tumors. Identifying the etiology of this failure has become a central focus of the field of cellular immunotherapy. Many pre-clinical studies have demonstrated that CAR engagement with antigen can rapidly drive the development of dysfunctional T cell states, such as exhaustion, as a result of the robust signaling activated by these synthetic receptors^1–4^. In addition to antigen-driven failure, CAR T cell failure can also result from antigen-independent “tonic” signaling, a feature that seems to be inherent to CAR structure^5,6^. Collectively, current data suggest that intrinsic to simplicity of CAR design is a predisposition to inducing T cell failure.

The native T cell receptor (TCR) is a large protein complex supported by a multitude of secondary supporting receptors, enabling exquisitely sensitive receptor triggering that orchestrates precise regulation of intracellular signaling quantity and quality. CARs, in contrast, are empowered with polyfunctionality but are deprived of the same precision control^7^. Recent evidence that TCR-based cell therapies can mediate meaningful responses in patients^8–10^ has re-invigorated efforts to harness TCR-driven immunity against cancer. Several novel approaches aim to bridge the benefits of MHC-independence of CAR and antibody-like antigen recognition using single chain variable fragment (scFv) adapters that link the TCR complex to cancer antigens^11–13^. Harkening back to original hybrid antigen receptor designs^14,15^, some groups have engineered novel antigen receptors by directly exchanging the TCR antigen binding domain with the antigen binding domain from an antibody.^16,17^ TCR and TCR-like cell therapies are, thus, emerging as an additional platform with the potential to enable native-like cellular immunity.

In parallel to others, we also designed hybrid antibody-TCR receptors, which we term an MHC-independent TCRs (miTCRs), using the antigen binding domain from the anti-CD19 antibody FMC63. We found these receptors to be highly functional but observed that a minor change in receptor structural orientation resulted in large distinctions in miTCR T cell functionality. This led us to interrogate the structure of this hybrid receptor. While CARs have benefitted from a resilience to domain hybridization, a fundamental principle of synthetic protein biology is that simple “cut” and “paste” of protein domains often does not result in predicted structure and function. Consistent with this, predictive modeling suggested that miTCR hybrid interfaces contained many biochemical conflicts. Through a series of prediction-informed alterations to miTCR structure and *in vitro* functional screens, we identified a variant miTCR that was predicted to have resolution of interface conflicts and that we confirmed drove significantly enhanced functionality. We further observed that, with costimulatory support, this variant miTCR was more durable and persistent *in vitro* and *in vivo* activity against large leukemic burdens than a classical CAR. Together, these studies use structure-informed protein design to develop a novel synthetic antigen receptor with enhanced anti-tumor function.

## Results

### Structural orientation of miTCRs impacts function

The TCR complex is a grouping of α and β chains, which are responsible for antigen binding, with CD3ο, ॐ, γ and ζ chains, responsible for intracellular signal transduction. Structurally, α and β chains are composed of constant regions (C_α_ and C_β_), which are the same across all receptors, and variable regions (V_α_ and V_β_), which are unique to each T cell clone and endow precise antigen-specificity. Antibodies have similar structural orientation, with constant and variable domains contained within their heavy and light chains and, like the TCR, the paired variable regions (V_H_ and V_L_) are responsible for antigen binding. Comparison of resolved protein structures^18,19^ confirmed that the extracellular TCR α and β chains are structurally similar to the antibody heavy and light chains (**Figure 1a**). This led us to hypothesize that exchange of these domains would lead to functional hybrid receptors that are MHC-independent and simultaneously engage native TCR regulatory circuits. To accomplish this, we replaced endogenous V_α_ and V_β_ sequences with the V_H_ and V_L_ sequences derived from the FMC63 antibody targeting CD19 (**Figure 1b**). Unlike other reports of similar receptors^16,17^, we developed both orientations of MHC-independent TCR (miTCR) design, V_L_/C_α_+V_H_/C_β_ (miTCR1) or V_H_/C_α_+V_L_/C_β_ (miTCR2). When expressed in a Jurkat reporter cell line, both miTCR1 and miTCR2 demonstrated antigen-specific activation of T cell transcription factors NFAT and NFκB, comparable to CD28 and 41BB-based CARs (**Supplementary figure 1a**) and confirming receptor functionality.

**Figure 1.**
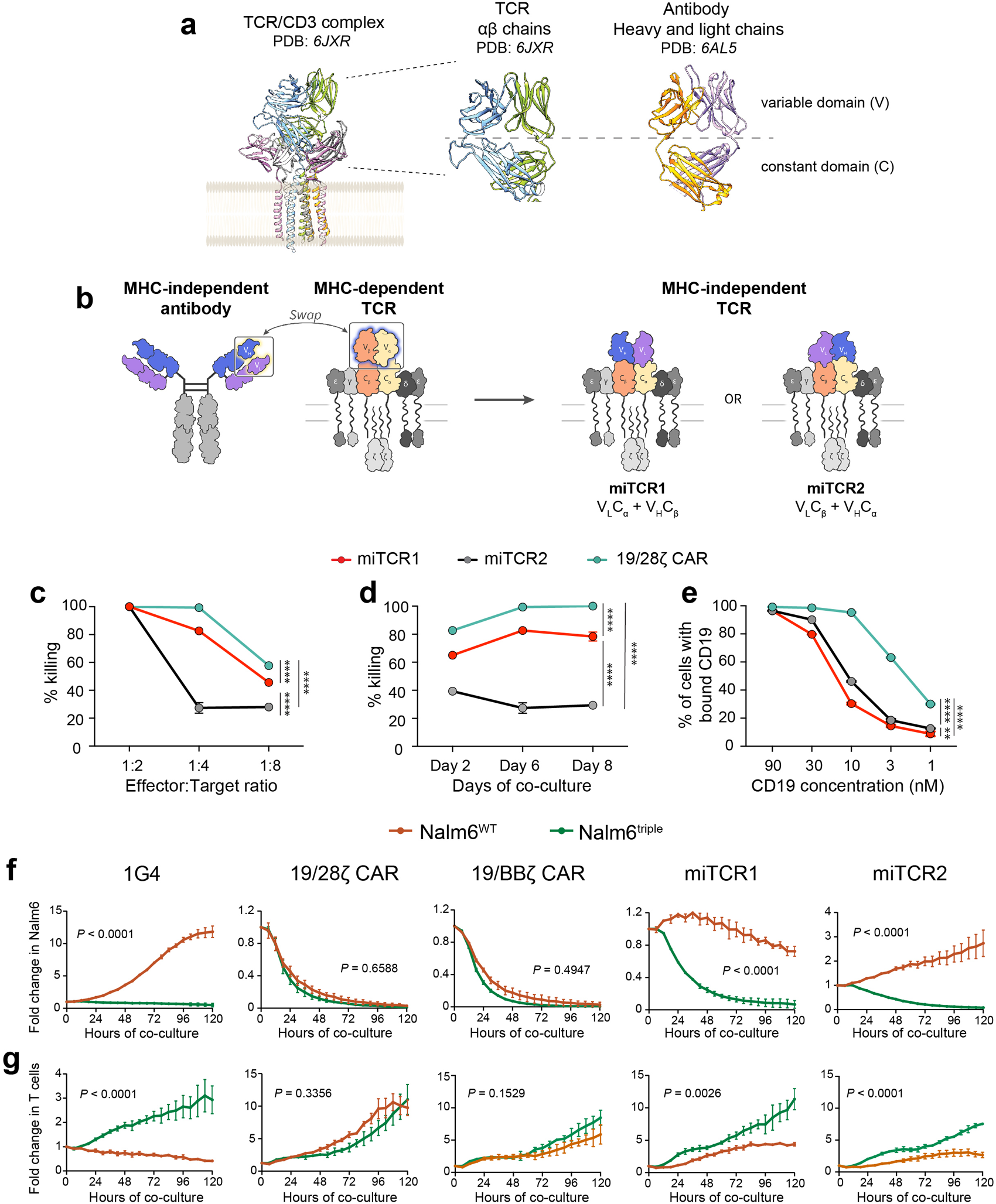
MHC-independent TCRs enable anti-tumor function. **a**, Comparison of TCR (cryo-EM) and antibody (crystal) variable and constant chain structures. **b,** Design of MHC-independent TCRs by swapping TCR variable regions with antibody variable regions to create two receptor formats. **c-d,** Cytotoxicity of miTCR1, miTCR2 and 19/28ζ CAR T cells against CD19+ Nalm6 cells **c,** after 6 days of co-culture at various E:T ratios and **d,** over time at an E:T ratio of 1:4. **e,** Percent of T cells that bound CD19-APC at various CD19 concentrations. **f,** Cytotoxicity, as measured by fold change in Nalm6 growth during co-culture, of each engineered T cell product against Nalm6^WT^ or Nalm6^triple^. **g,** T cell expansion, as measured by fold change in engineered T cells during co-culture, of each engineered T cell product against Nalm6^WT^ or Nalm6^triple^. **c-g** are representative data of five independent donors. \*\**P <* 0.01, \*\*\**P <* 0.001, \*\*\*\**P* < 0.0001 by two-way ANOVA (**c-e**) or Student’s t-test (**f-g**).

To prevent mispairing of miTCR chains with endogenous TCR chains in primary human T cells, we disrupted both *TRAC* and *TRBC* genes using CRISPR/Cas9 genome editing, leading to nearly 100% loss of surface TCR (**Supplementary figures 1b-c**). Using this platform, miTCR expression could reliably be confirmed by evaluating surface TCR constant chains or the variable region of FMC63 (**Supplementary figures 2a-b**). Indeed, co-staining for both demonstrated a 1:1 correlation, confirming that all surface TCRs were miTCRs (**Supplementary figure 2c**). Further, expression of miTCRs resulted in returned surface expression of CD3ε, which requires pairing with TCR constant chains prior to membrane integration, establishing that miTCRs associate with native TCR machinery (**Supplementary figure 2d**). We observed that despite similar transduction efficiency (as assessed by expression of an mCherry marker expressed on the miTCR construct), miTCR2 surface expression was consistently higher than miTCR1 (**Supplementary figure 2e**). Paradoxically, however, *in vitro* cytotoxicity assays normalized to mCherry+ T cell effectors revealed that miTCR1 T cells were more effective at killing CD19+ human B cell leukemia (Nalm6) despite this lower receptor expression (**Figures 1c-d**). We further found that while both miTCRs had lower CD19 binding capacity than a standard CD19 CAR, miTCR1 binding capacity was significantly lower than miTCR2 (**Figure 1e, Supplementary figure 2f**).

Native TCRs require support from costimulatory receptors to enable full T cell function. To test the impact of costimulation on miTCR function we engineered Nalm6 cells to express 41BBL, CD80 and CD86 (Nalm6^triple^), the ligands for the costimulatory receptors that are integrated into current CAR design (41BB and CD28). In parallel, we engineered T cells to express one of two CD19-targeted CARs (19/28ζ or 19/BBζ) or a transgenic TCR targeting the cancer-associated antigen NYESO1_157-165_ in HLA-A*02:01 (TCR clone 1G4^20^). To enable recognition of Nalm6 by 1G4 T cells, we further engineered Nalm6 cells to express HLA-A*02:01 and full-length NYESO1. Predictably, 1G4 T cell anti-tumor cytotoxicity (**Figure 1f**) and expansion (**Figure 1g**) were significantly improved when combined with Nalm6^triple^, while additional costimulation had minimal impact on 19/28ζ or 19/BBζ CAR T cell function. Like their TCR counterparts, miTCR1 and miTCR2 T cells had significantly greater function when targeting Nalm6^triple^. Intriguingly, miTCR1 appeared less reliant on costimulation than miTCR2, demonstrating reasonable cytotoxic function against Nalm6^WT^. Direct comparison of receptor functionality revealed that in the presence and absence of costimulation, miTCR cytotoxicity and expansion was broadly more similar to CARs than TCRs (**Supplementary figure 3a-d**). Directed comparison again revealed enhanced cytotoxicity and T cell expansion driven by miTCR1, confirming a distinction in function that was dependent on orientation of domain hybridization.

### miTCR design results in biochemical conflicts at the variable-constant domain interface

We were motivated to understand why two receptors composed of the same four core domains differed in surface expression and function. While all components of miTCR structure are naturally-expressed extracellular proteins, the interface created by linking antibody variable and TCR constant regions is synthetic (**Figure 2a**). The molecular structures of our miTCRs, or other similar receptors, have not been resolved and thus we relied on predictive modeling, a field that has seen tremendous progress in reliability^21,22^, to interrogate miTCR structure.

**Figure 2.**
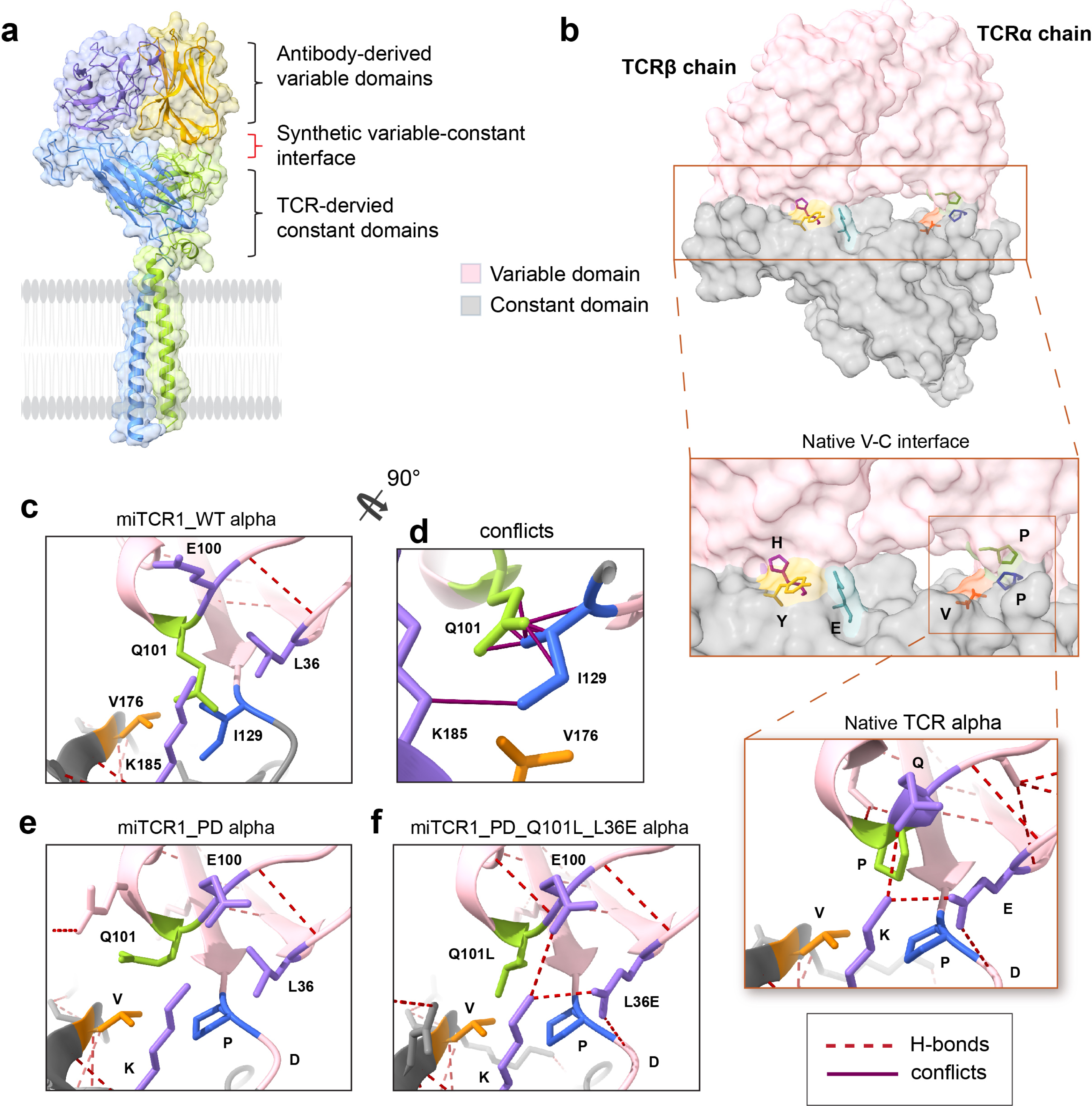
miTCR V-C interface contains inherent biochemical conflicts. **a**, Predictive modeling of miTCR1 paired α and β chains. **b,** Detailed analysis of cryo-EM resolved variable-constant (V-C) interface of the native TCR (PDB: *6JXR*). **c,** Predictive modeling of the miTCR1α V-C interface. **d,** Rotation of the V-C interface to reveal predicted biochemical and structural conflicts. **e,** Predictive modeling of the miTCR1α V-C interface with PD insertion **f,** and Q101L and L36E mutations. Modeling performed using Phyre2 or AlphaFold 2.

The overall structures of miTCR1 and miTCR2 were predicted to be very similar to the cryo electron microscopy (cryo-EM) resolved TCRα and β chain structures^18^ (**Supplementary figure 4a**), with 86-91% of each miTCR chain mapping to the native TCR chains with 100% confidence. Interrogation of the TCRα chain variable-constant (V-C) interface revealed highly-ordered structure pivoting around a hydrophobic triangle anchored by two prolines and a valine (P-V-P), which was itself surrounded by a hydrophilic cage of glutamine, glutamic acid and lysine (Q-E-K, **Figure 2b**). Overlay of the five top ranking predicted V-C interface structures of miTCR1 and miTCR2 demonstrated exquisite consistency between models, providing high confidence in the reliability of predicted miTCR V-C interfaces (**Supplementary figure 4b**). In contrast to the TCR V-C interface, the miTCR1α V-C interface was predicted to be disrupted, resulting from intrusion of an isoleucine (I129) and charged glutamic acid (Q101) from the V_L_ domain into the hydrophobic core (**Figure 2c**). These changes were predicted to cause significant biochemical conflicts, partially resulting from close proximity of Q101 to I129, predicted to disrupt stabilizing hydrogen bonds and cause torsion of the V_L_ and C_α_ interface (**Figure 2d**). Similarly, miTCR2α was also predicted to have disruption of its hydrophobic core (**Supplementary figure 4c**). TCRβ also has a hydrophobic core at the V-C interface (**Supplementary figure 4d**), and predictive modeling of miTCR1β and miTCR2β chains identified changes in the same two interface amino acids (K and F) but with different changes (miTCR1β K>G, F>P; miTCR2β K>S, F>S, **Supplementary figure 4d**).

Based on these observed deviations from native TCR structure, we hypothesized that the synthetic V-C interface structure had a central role on miTCR expression and function. To evaluate this, we performed iterative amino acid alterations and predictive modeling with the goal of alleviating V-C interface conflicts. To illustrate one example of alterations made to the miTCR1α chain, we first introduced native proline-aspartic acid (PD) to resolve steric disruption from I129 (**Figure 2e**). While this was predicted to improve some strain, the intruding Q101 continued to disrupt the hydrophobic core. We mutated Q101 to a leucine and neighboring L36 to a glutamic acid, which enabled recreation of the hydrophobic core triangle (now P-V-L instead of P-V-P), surrounding hydrophilic cage (now E-E-K instead of Q-E-K) and supportive hydrogen bonds (**Figure 2f**). Through similar processes, we generated a series of variants with mutations, insertions and/or deletions at the V-C interface for all four chains (representative examples for each chain illustrated in **Supplementary figure 4d**) resulting a panel of 10 miTCR1 and 6 miTCR2 structures (**Supplementary figure 5**).

### Interface alterations improve function of miTCRs

We next undertook a series of *in vitro* studies to elucidate the functional implications of these interface alterations. Most miTCR1 variants (mut031, 032, 034, 035, 037) and all miTCR2 variants compelled improved surface receptor expression, with some (mut040, 042 and 044) demonstrating comparable expression to the natively-structured 1G4 when expressed in Jurkats (**Figure 3a**) and human T cells (**Supplementary figure 6a**). Like their wild-type counterparts, miTCR2 variants consistently demonstrated higher surface expression (**Supplementary figures 6b-c**). By virtue of our iterative design, we were able to test the specific impact of individual changes and observed that, independently of any other changes made, addition of the PD domain into the miTCRα chain consistently improved variant surface expression (**Supplementary figure 6d**), highlighting the importance of this single alteration. Jurkat reporter cells engineered to express these variants demonstrated potent and antigen-specific activation of NFAT and NFκB in response to CD19, confirming functionality of all variant receptors (**Supplementary figures 6e-f**). Armed with this panel of variants we performed a functional *in vitro* screen in human T cells to test the impact of interface alterations on receptor-driven cytotoxicity and T cell expansion, using the potent 19/28ζ CAR as a control. In co-cultures against Nalm6^WT^, miTCR1 variants broadly outperformed miTCR2 variants (**Figure 3b**). The addition of costimulation again improved miTCR-driven T cell functionality, with a recurrent demonstration that miTCR2-like structures were more reliant on costimulation (**Figure 3c**). T cell expansion mirrored cytotoxic function, with miTCR1 variants outperforming miTCR2 variant T cells but inferior to CAR T cells without costimulation (**Supplementary figure 6g**). With costimulation, however, many miTCRs induced prolonged T cell expansion even after tumor clearance, with some miTCR1 variant T cells expanding better than 19/28ζ CAR T cells (**Supplementary figure 6h**).

**Figure 3.**
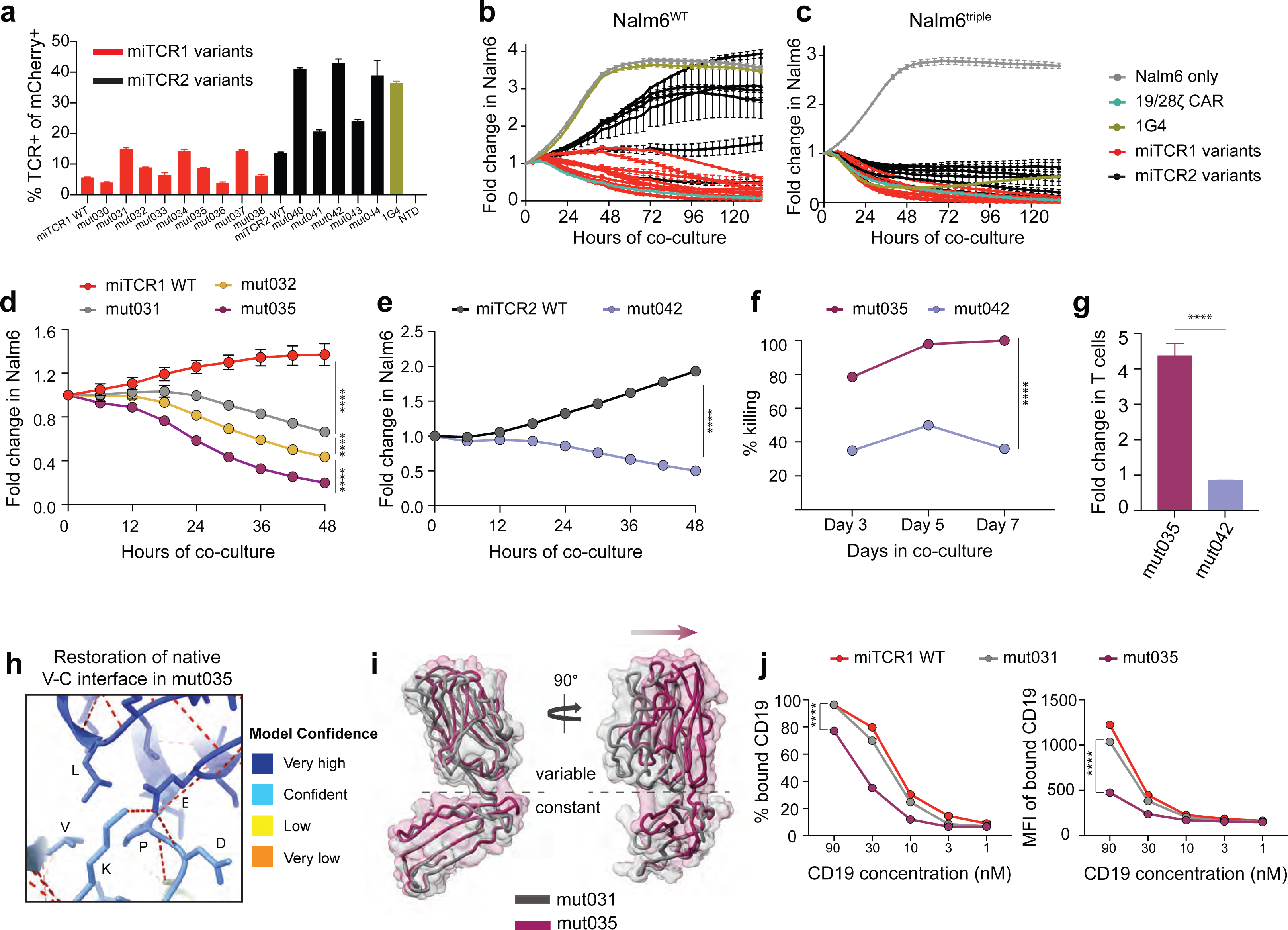
Targeted V-C interface alterations improve miTCR function. **a**, Expression of TCR constant chains on the surface of Jurkats transduced with miTCR variants. **b-c,** Functional screen of miTCR variant-expressing human T cells to evaluate cytotoxicity against **b,** Nalm6^WT^ or **c,** Nalm6^triple^. **d-e,** Cytotoxicity of lead **d,** miTCR1 and **e,** miTCR2 variants against Nalm6^WT^ at an E:T of 1:4. **f,** Cytotoxicity of miTCR1 mut035 and mut042 against high burden Nalm6^WT^ (E:T 1:8) over time. **g,** T cell expansion of mut035 and mut042 at the conclusion of 7 day co-cultures (E:T 1:8). **h,** Predictive modeling of the V-C interface of mut035. **i,** Predictive modeling and overlay of mut031 and mut035 α chains. **j,** CD19 binding capacity of miTCR1 WT, mut031 and mut035 measured by percent of T cells that bound CD19-APC at various CD19 concentrations and MFI of bound CD19. Screening studies in **b-c** performed using two independent donors. **d-g** are representative data from three independent donors. Modeling in **h-i** performed using AlphaFold 2. \**P <* 0.05, \*\**P <* 0.01, \*\*\**P <* 0.001, \*\*\*\**P* < 0.0001 by two-way ANOVA (**d, j**) or by Student’s t-test (**e-g**).

From this screen, we aimed to validate the most effective candidate receptors. Dedicated cytotoxicity studies confirmed that mut031, 032 and 035 were the most effective miTCR1 variants, with mut035 leading in functionality (**Figure 3d, Supplementary figure 7a**), and that mut042 was the most effective miTCR2 variant (**Figure 3e, Supplementary figure 7b**). This hierarchy of function was further confirmed using distinct but complementary long-term flow cytometry assays (**Supplementary figures 7c-d**). We next tested mut035 and mut042 in long-term *in vitro* stress assays with higher leukemic burden (1 T cell per 8 Nalm6^WT^). These studies clearly demonstrated that mut035 was superior to mut042, driving more cytotoxicity against Nalm6^WT^ and greater T cell proliferation (**Figures 3f-g**).

Having identified the most potent miTCR variant, we aimed to understand how its structure differed from other similar variants. Mut031 and mut035 differed only at two residues: both contained the PD insertion mentioned above, but mut031 retained the FMC63 V_L_ residues Q101 and L36 while mut035 had been mutated at these sites (Q101>L and L36>E). We compared the predicted mut031 V_L_ structure to the recently resolved^23^ cryo-EM structure of FMC63 V_L_ and found them to be nearly identical (**Supplementary figure 7e**). Given the high predictive confidence for this protein region, we then modeled the mut035 V-C interface to find that Q101L and L36E enabled restoration of the native TCR V-C interface with very high predictive confidence (**Figure 3h**). Notably, this modeling was performed using a distinct system (AlphaFold 2^22^) than our iterative mutation/prediction modeling in Figure 2 (Phyre2^21^), adding further confidence to the resolution of interface conflicts in mut035. Modeling of the entire mut031 and mut035 α chains predicted that these two mutations not only caused changes at the V-C interface but also a shift of the distal antigen binding domains (**Figure 3i**), which was associated with a reduction in CD19 binding capacity (**Figure 3j**). Collectively, these data indicate that targeted alterations to the V-C interface can significantly improve miTCR-driven function.

### miTCR T cells outperform CAR T cells against high burdens of leukemia

We found that while there was variability in the degree of reliance on costimulation, all miTCRs benefited from costimulatory support. Our previous studies engaged both 41BB and CD28 pathways, as have other miTCR-like products^17^. To more fairly compare miTCR function to that of commercially-available CAR products, which are supported by only one costimulatory domain, we specifically aimed to engage only one pathway. Mut035 T cells were co-cultured with Nalm6 cells engineered to express 41BBL (Nalm6^41BBL^), CD80/86 (Nalm6^CD80/CD86^) or all three as in previous studies (Nalm6^triple^). Importantly, all cell lines had equivalent growth kinetics (**Supplementary figure 8a**). We found that mut035 T cells were able to effectively eliminate each Nalm6 cell line with similar kinetics in these acute cytotoxicity assays (**Figure 4a**). Evaluation of mut035 T cell expansion, however, revealed dynamics similar to what has been shown for CAR T cells: CD28 support increased the quantity of expansion while 41BB supported prolonged duration of expansion (**Figure 4b**). Given this similarity in cytotoxicity, we undertook re-challenge studies to stress the mut035-driven T cell response. Following clearance of initial Nalm6 burdens and 2-3 days of leukemia-free rest, we re-challenged mut035 T cells with Nalm6^WT^ to determine how initial costimulation impacted a secondary response. We observed that initial exposure to either Nalm6^41BBL^ or Nalm6^triple^ allowed robust clearance of this second tumor challenge, while initial exposure to Nalm6^CD80/86^ led to a hypofunctional response (**Figure 4c**). While overall mut035 T cell expansion in response to Nalm6^WT^ was modest, it was not impacted by initial stimulation (**Figure 4d**), highlighting that these differences in function were qualitative and not a result of differences in T cell quantity.

**Figure 4.**
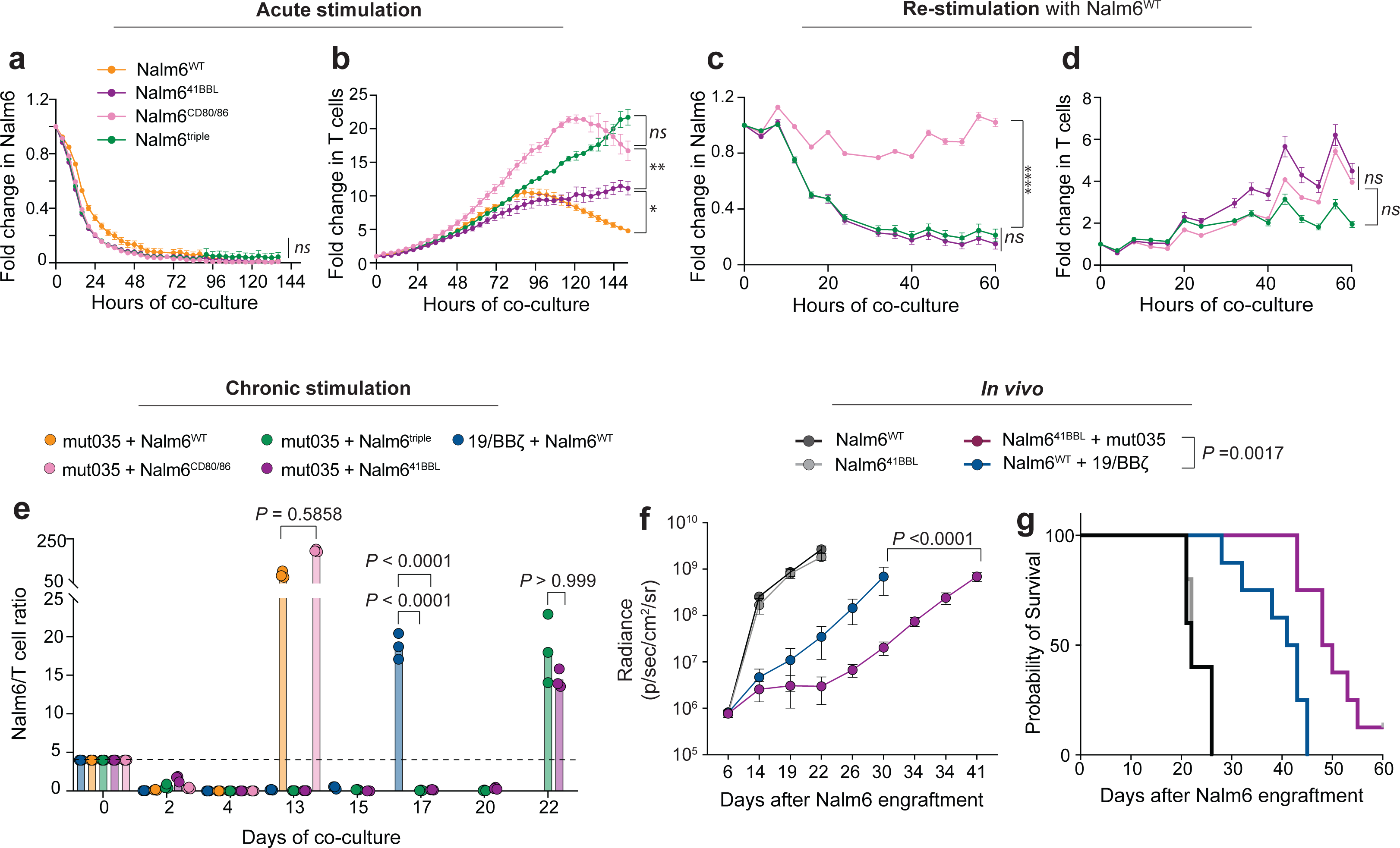
mut035 enables enhanced durability against high burdens of leukemia. **a**, **Cytotoxicity and b,** T cell expansion of mut035 T cells against various Nalm6 targets. **c,** Cytotoxicity and **d,** T cell expansion of mut035 T cells upon re-challenge with Nalm6^WT^ targets. **e,** Nalm6 cells per T cell over time during chronic stimulation co-cultures. **f-g,** Change in *in vivo* disease burden over time as measured by bioluminescent signal and **g,** survival of NSG mice engrafted with either Nalm6^WT^ or Nalm6^41BBL^ and treated with low-dose (2.5x10^5^) T cells. For **a-e,** representative data from three independent donors. \**P <* 0.05, \*\**P <* 0.01, \*\*\**P <* 0.001, \*\*\*\**P* < 0.0001 by two-way ANOVA. For **f-g,** n=8 animals per group; significance determined using two-way ANOVA (**a-f**) or Log-Rank test (**g**).

Based on these data, we selected 41BB as the optimal single costimulatory support signal for miTCR T cells and compared mut035 T cells to 19/BBζ CAR T cells. To model equivalent costimulatory support, we combined mut035 T cells with Nalm6^41BBL^ and 19/BBζ CAR T cells against Nalm6^WT^, providing a “single” costimulatory signal to each. Both acute and re-challenge studies demonstrated equivalent cytotoxicity between 19/BBζ CAR T cells and mut035 T cells (**Supplementary figures 8b-c**), and thus we moved to a more stressful *in vitro* model. We previously developed a protocol that reliably induces 19/BBζ CAR T cell dysfunction following 12-17 days of chronic antigen stimulation^2,24^. We established co-cultures of 19/BBζ CAR T cells with Nalm6^WT^ and mut035 T cells with Nalm6^41BBL^, as well as Nalm6^WT^, Nalm6^CD80/86^ and Nalm6^triple^, and replenished these with fresh tumor every other day to maintain persistent stimulation. Measurement of surface receptor expression revealed that CARs retained robust surface expression during chronic stimulation, whereas mut035 miTCRs were down-regulated, a process that was stabilized by the presence of costimulation (**Supplementary figure 8d**). Chronic stimulation with Nalm6^WT^ or Nalm6^CD80/CD86^ demonstrated poor mut035 expansion (**Supplementary figure 8e**) and early loss of tumor control at ∼day 13 of chronic stimulation (**Figures 4e**). Consistent with our previous findings^2^, 19/BBζ CAR T cells expanded and contracted rapidly, losing tumor control ∼day 17. Chronic stimulation of mut035 T cells with Nalm6^41BBL^ or Nalm6^triple^, however, resulted in a more modest but more durable T cell expansion and a ∼9-day delay in T cell contraction compared to CAR T cells (**Supplementary figure 8e**). Consistently, mut035 T cells demonstrated more durable control of Nalm6^41BBL^ and Nalm6^triple^, losing control ∼day 22 (**Figure 4e**). To further test the durability of mut035 T cell functionality, we employed an *in vivo* stress model in which immunodeficient (NOD/SCID/γc^-/-^) mice were engrafted with Nalm6^WT^ or Nalm6^41BBL^ and given a low dose (2.5x10^5^) of engineered T cells seven days later^2,25^. Tracking of disease using bioluminescent imaging confirmed identical growth kinetics of these two tumors *in vivo* and demonstrated that mut035 T cells significantly delayed disease progression as compared to 19/BBζ CAR T cells (**Figure 4f**). As expected, no animals were cured in this stress model but, consistent with enhanced disease control, mut035 T cells significantly prolonged animal survival (**Figure 4g**, *P = 0.0017*). Collectively, these data demonstrate that, with 41BB costimulation, mut035 miTCR T cells exert more durable control of large leukemic burdens than 19/BBζ CAR T cells.

## Discussion

The widespread and cost-effective application of gene engineering and genome editing now permits rapid and iterative evaluation of synthetic proteins in primary human cells. Of the many translational applications, the paradigm of synthetic biology is the chimeric antigen receptor. Despite the breakthrough success of CD19-directed CAR therapies, these products eventually fail to control disease in >50% of patients. No other CAR-based platforms have recreated the success of CD19 CARs and, as a result, myriad efforts from both academic and industry researchers aim to capitalize on the proof that synthetic proteins can redirect T cell function but instead engineer native TCR-driven cell function. Amongst the many remarkable features of CAR design is its resilience to linking unrelated protein domains. Here, we demonstrate that domain hybridization is more complex for TCR-based hybrid receptor formats, underscoring the central roles of biochemistry and structural biology in synthetic immunotherapy. We identify an approach to overcome structural complexity using protein modeling and iterative receptor engineering to meaningfully enhance hybrid receptor function against large tumor burdens.

While experimental resolution of protein molecular structure remains important for many applications, recent improvements in accuracy of predictive modeling platforms enable structural interrogation without the requirement of complex and labor-intensive studies. Using these advanced platforms, we identified alterations in amino acids that were predicted to stabilize the V-C interface of miTCRs and experimentally confirmed them to enhance functionality. One notable observation is that all variants improved function compared to wild-type miTCRs, except for two (mut037 and mut038, which were equivalent to WT), underscoring the need to experimentally test the impact of structural changes. Broadly, however, stabilization of the V-C interface was beneficial to miTCR-driven T cell function. Another notable observation is that miTCR surface expression, either of wild-type or variant receptors, did not correlate with function. This questions the presumption that “more is better” when manufacturing engineered cell therapies and suggests instead a more complex structure/function relationship for miTCRs. While this report focuses on CD19-targeting using FMC63, we have also explored several other miTCR designs targeting distinct antigens expressed by hematologic and solid tumors. Not surprisingly, each variable-constant pairing brings unique challenges to interface structure, and we are actively exploring the functional implications of altering these structures in several contexts.

Analysis of antigen receptor structure is often focused on the antigen binding domains, specifically the complementarity determining regions (CDRs) that regulate antigen recognition^26^, however increasing attention is now being paid to the framework regions that support these distal domains. One recent study used predictive modeling of the framework of native TCR variable chains to identify specific residues that regulated surface TCR expression and receptor-driven function^27^. These observations suggest that in addition to CDR variability, naturally evolved structural variation within TCR framework plays a role in controlling endogenous immune responses. Our data build on these observations to demonstrate the importance of miTCR framework residues in development of synthetic antigen-specific immunity.

We are not the first to evaluate MHC-independent TCR formats. T cell antigen couplers (TACs)^11^ and T cell receptor fusion constructs (TRuCs)^12^ use single chain variable fragments to target the native TCR to cell surface antigens, but neither of these alter TCR structure. The concept of domain hybridization to generate an MHC-independent TCR is among the original chimeric receptor designs, dating to a 1989 report from Esshar and colleagues^14^. This approach was not heavily re-investigated until recently, in part due to engineering barriers such as construct size and the need to prevent mispairing with endogenous TCR chains, now easily surmountable. Synthetic T cell and antigen receptors (STAR)^16^ and HLA-independent TCRs (HIT)^17^ are similar in design to our wild-type miTCRs and have recently been shown to mediate potent anti-tumor activity. Our data add to the growing body of literature related to this receptor format and make several complementary and supportive observations, among which is the role of costimulation. Recognizing that our method of costimulation is highly artificial, we identified that, like HITs, miTCRs are less reliant on costimulation than classical TCRs. While the biology underlying this phenomenon is an area of ongoing investigation, we speculate that this may be a result of differences in affinity between V_L_/V_H_ and V_α_/V_β_. Several studies have revealed a complex relationship between affinity and receptor-driven T cell function. Long-standing data of TCRs^28^ and recent data of CARs^29^ have suggested that very high affinities are detrimental to T cell function, consistent with our observed association between function and antigen binding capacity of miTCRs. Intriguingly, we observed that mut035 T cells respond similarly to Nalm6^41BBL^ and Nalm6^triple^, while responses to Nalm6^CD80/86^ were inferior. Recent studies of CAR T cells have demonstrated synergy when 41BB and CD28 are delivered in *trans*^30–32^, and dissecting differences in the impact of synthetic and endogenous costimulation on CARs and miTCRs is critical to the design of novel cellular therapies.

While alike in concept and several key observations, our studies differ from the previous STAR and HIT reports in important ways. First, we evaluated a multitude of receptor structures, permitting us to “evolve” a more functional receptor architecture using *in vitro* screening. Second, these platforms utilized distinct methods to prevent mispairing of introduced transgenic α and β chains with endogenous chains; STARs incorporate potentially immunogenic murine TCR constant regions, complicating clinical translation, while HITs use homology-directed repair (HDR) strategies to directly insert their receptor into the *TRAC* locus, preventing mispairing of endogenous TCRα with HITβ but permitting mispairing of TCRβ with HITα. We instead used dual disruption of *TRAC* and *TRBC* to abrogate expression of endogenous TCRs. Third, and perhaps most importantly, both STAR and HIT demonstrate their most potent efficacy, and superiority to CARs, in response to low-antigen tumors. Our initial design of wild-type miTCRs demonstrated inferior function compared to CD19-targeted CARs against CD19^hi^ ALL, but through structural modifications our lead variant (mut035) was superior not only to wild-type miTCRs but also CD19 CARs. Improving responses to low-antigen tumors is an important clinical barrier but is primarily relevant to patients who have already failed immunotherapy as a result of immune selection. For current cellular immunotherapy targets, such as CD19 and BCMA, disease at time of treatment is almost always antigen “high”. CD19 loss is often observed in children and young adults who relapse after treatment for ALL with a CD19-directed therapy, although antigen “low” relapses are less common^33^. Data from CD22 CAR clinical trials suggest that CD22 downregulation, but not loss, may be a significant cause of escape, identifying a clear need for therapies that can re-target CD22^low^ disease in this setting^34^. Beyond treatment of relapse, targeting low antigen diseases is complicated by the increased potential for off-tumor on-target toxicity that could be mediated by a receptor with high sensitivity for low antigen quantity. This is particularly relevant in solid tumors, in which cancer-specific antigens are rare and antigen-targeted therapies often rely on increased expression of target antigens by cancer cells as compared to healthy counterparts. We thus believe that a specific advantage of our approach is the ability to target high density antigens and simultaneously enhance T cell function.

One aspect of TCR-driven biology that was not addressed in this study was the impact of miTCRs on immunotherapy toxicities. Cytokine release syndrome (CRS) has been shown to be driven by excessive CAR-driven T cell activation that stimulates pathologic IL6 production by myeloid-lineage cells^35–37^. An area of ongoing investigation is how miTCR T cells engage endogenous innate immune cells, and the downstream impact on CRS. While the biology is not as clear, the impact of a TCR-like receptor on the development of immune cell neurotoxicities, which have several varieties^38,39^, is also of significant interest.

In summary, we report that in-depth interrogation of predicted structures of hybrid receptors can enable rational synthetic interventions that improve T cell function.

## Acknowledgements

We thank Dr. Carl June (University of Pennsylvania) for generously providing the 1G4 TCR construct and Dr. Peter Steinberger (Medical University of Vienna) for providing the Jurkat Triple Reporter Cell lines. These studies were supported by the Damon Runyoun Foundation Clinical Investigator Award, K08CA237740, and the Be The Match Foundation Amy Strelzer Manasevit Award (all to N.S.).

## Author contributions

J.F.-C. and N.S. designed and oversaw the research. J.H.L., T.-C.C., Y.T., J.M.W., M.E.S., J.R., Y.-S.H., M.S., D.G. and J.F.-C. performed the research. J.F.D. provided technical and conceptual support. J.F.-C. and N.S. wrote the manuscript. All authors reviewed the manuscript.

## Competing interests

J.-F.C. and N.S. have submitted patent applications related to this work. N.S. is an inventor on patents related to adoptive cell therapies, held by Washington University and the University of Pennsylvania (some licensed to Novartis). Unrelated to this work, N.S. has served as a consultant for several companies involved in cell therapies and is a board member for Phoreus Biotech. J.F.D. receives research funding from Amphivena Therapeutics, NeoImmuneTech, Macrogenics, Incyte, Bioline Rx, Wugen; has equity ownership in Magenta Therapeutics, Wugen; consults for Incyte, RiverVest Venture Partners, hC Bioscience, Inc.; and is a board member for RiverVest Venture Partners, Magenta Therapeutics.

## Data Availability

All requests for data will be reviewed promptly by the corresponding author and made available, pending any intellectual property obligations. Any materials that can be shared will be via a material transfer agreement.

## Methods

### General cell culture and flow cytometry

Cells were grown and cultured at a concentration of 1x10^6^ cells/mL of standard R10 culture media (RPMI 1640 + 10% FBS, 1% penicillin/streptomycin, 1% HEPES, 1% glutamine) at 37°C in 5% ambient CO_2_. Samples were stained with antibodies against TCR constant chains (Biolegend, clone IP26, #306741), CD3ε (BD Horizon, clone UCHT1, #563109), FMC63 scFv (Acro biosystems, Y45, #FM3-BY54) in 100ul FACS buffer (2% FBS in PBS), washed once with the same buffer and analyzed on the Attune NxT Flow Cytometer (ThermoFisher). Cells were gated and analyzed using FlowJo v9 or 10 (BD Biosciences).

### Vector construction

All constructs were generated by inserting synthesized and codon optimized target sequences (GenScript) into a lentiviral transfer plasmid backbone containing an EF1α promoter to drive target gene expression. The leader sequence for both alpha chain (METLLGLLILWLQLQWVSSKQ) and beta chain (MSIGLLCCAALSLLWAGPVNAGV) were derived from 1G4, a transgenic TCR targeting HLA-A2 restricted NY-ESO-1^20^. The variable light and heavy chain sequences of miTCR were derived from anti-CD19 antibody clone FMC63. The human TCR α and β constant region sequences were derived from published sequences (UniProtKB P01848 and A0A5B9, respectively). A cysteine mutation was introduced into each constant chain to promote interchain disulfide bonding for optimal TCR pairing^40^. The two TCR chains were separated by a furin-GSG-T2A peptide^41^ and followed by a GSG-P2A linked to an mCherry transduction marker.

### Lentiviral engineering and transduction of human T cells

Lentiviral vectors were manufactured as previously described^42^. Briefly, pMDG.1 (7 μg), pRSV.rev (18 μg), pMDLg/p.RRE (18 μg) packaging plasmids and 15 μg of expression plasmids were mixed and transfected into 293T cells using Lipofectamine 3000 (Invitrogen #L3000150) according to manufacturer’s protocol. At both 24 and 48 h following transfection, supernatant was collected and filtered through 0.45μm aPES filters (Thermo scientific #1650045). Virus containing media was then concentrated using high-speed centrifugation (8500×*g,* 16-18hr at 4°C with attenuated deceleration). Virus particles were re-suspended in 293T growth medium (1/100∼1/200 of the original volume) and snap-frozen. The virus was stored at -80°C before usage. For T cell engineering, PBMCs were procured from leukoreduction chambers, and CD4 and CD8 cells were purified using magnetic beads (Miltenyi Biotec) and combined at a 1:1 ratio before freezing. T cells were activated using CD3/CD28 stimulatory beads (DynaBeads Thermo-Fisher #40203D) at a ratio of 3 beads/cell and incubated at 37°C. 24-30 hr after stimulation, lentiviral vectors encoding miTCRs, transgenic TCRs or CARs were added to stimulatory T cell/beads cultures at a MOI of 2-4. Beads were removed after 4-6 days of stimulation. Edited cells were cultured and expanded in R10 containing 5ng/ml IL7 and 5ng/ml IL15 (Peprotech #20007, #20015) and replenished every 2-3 days until freezing at days 14-16.

### Phyre2 and AlphaFold 2 structural modeling

Three-dimentional structures were predicted using Phyre2^21^ or AlphaFold 2^22^. The resulting models containing either wild-type or mutated chains were superimposed with native TCR structure (PDB: 6JXR) and visualized by UCSF ChimeraX^43^.

### CRISPR editing

sgRNAs for *TRAC* (TGTGCTAGACATGAGGTCTA) and *TRBC* (GGAGAATGACGAGTGGACCC) were based on previously published sequences^44^ and purchased from IDT. Ribonucleoprotein (RNP) complexes were formed by incubating 10µg of TrueCut™ Cas9 Protein v2 (Invitrogen #A36499) with 20µg of each sgRNA for 10 min at room temperature. T cells, either resting or activated, were washed once with room temperature PBS (Gibco #14190136) and spun at 200xg for 10 min and resuspended at a concentration of 2-10x10^6^ cells/100µL in Lonza Buffer P3/supplement (Lonza #V4XP-3024). The RNP complex and 100µL of resuspended cells were combined and electroporated using pulse code EO-115 on the Lonza 4D-Nucleofector Core/X Unit.

### Jurkat cell transfection and activity assays

1-1.5 x10^6^ triple reporter TCR^-/-^ Jurkat cells were mixed with 2μg of pLVM containing gene of interest and electroporated using pulse code CL-120 according to manufacturer’s protocol (Lonza #V4XC-1024, SE Cell line kit). After 48-72hrs, miTCR or CAR expression on cells was analyzed by flow cytometry. Jurkat cells were combined at 1:1 ratios with either Nalm6 (CD19+) or Molm14 (CD19-) cells and incubated at 37°C overnight and analyzed by flow cytometry the following day to assess transcription factor activation.

### CD19 binding assays

CD19-APC was made by using APC conjugation kit (Abcam # ab201807). Briefly, 50ug of CD19 protein (Sino Biological #11880-H02H) was reconstituted in 90μl water and 10μl of modifier was added to the protein solution. The protein mixture was then mixed with the APC conjugation reagent overnight at room temperature. 11μl of quencher was added for 30min before usage. Primary human T cells expressing CAR or miTCRs were incubated in 100μl T cell growth medium R10 containing a gradient of increasing concentration of CD19-APC at 37°C for 30min on 96well plate. 1ul of anti-TCR antibody was then added to the well, and the incubation continued for another 30min at 37°C (total 1hr incubation time for T cells with CD19-APC). After washing twice with FACS buffer, cells were analyzed by flow cytometry.

### Acute, re-stimulation and chronic stimulation assays

GFP+ Nalm6 cells were co-cultured with T cells in technical triplicate at E:T ratios of 1:4 unless otherwise stated in 200ul T cell medium in 96-well flat bottom plates (Cellstar, #655180). Plates were imaged every 4-6 hours for 3-7 days using the IncuCyte*®* Live*-*Cell Analysis System (Sartorius). 5 images per well at 10x zoom were collected at each time point. Total GFP area (μm^2^/well) and total integrated RFP intensity or total RFP area per well were used as a quantitative measure of live cancer and T cells, respectively. Values were normalized to the time=0 measurement. For re-stimulation experiments, T cells were first harvested at the end of initial acute stimulation and re-quantified using flow cytometry, then re-combined with Nalm6 target cells at indicated E:T ratios. The cell mixtures were then recorded by IncuCyte as described above. For flow cytometry-based cytotoxicity assays, T cells were combined with target cells at E:T ratios of 1:4 unless otherwise stated, and co-cultures were evaluated for absolute count of target and T cells by flow cytometry. All co-cultures were established in technical triplicate.

Chronic stimulation assays were performed as previously described^24^. Briefly, absolute count of mCherry+ T cells and GFP+ Nalm6 were evaluated by flow cytometry and combined with an E:T ratio of 1:4. The absolute count of T cell and cancer cells in the mixture were re-evaluated every 2-3 days by flow cytometry. Fresh cancer cells were added to the mixture to main an E:T ratio of 1:4 until T cells failed to control cancer cell growth (E:T ratio exceeded 4).

### Xenograft mouse models

6-10 week old NOD-SCID-γc^-/-^ (NSG) mice were obtained from the Jackson Laboratory and maintained in pathogen-free conditions. Animals were injected via tail vein with 1x10^6^ Nalm6 WT or Nalm6^41BBL^ cells in 0.2mL sterile PBS. On day 7 after tumor delivery, 0.25x10^6^ CAR or miTCR+ T were injected via tail vein in 0.2mL sterile PBS. Animals were monitored for signs of disease progression and overt toxicity, such as xenogeneic graft-versus-host disease, as evidenced by >10% loss in body weight, loss of fur, diarrhea, conjunctivitis and disease-related hind limb paralysis. Disease burdens were monitored over time using Spectral Instruments Imaging AMI Instrument with analysis done using associated Aura software.

Animals were sacrificed when radiance reached >3x10^9^ photos/sec/cm2/sr (5-log greater than background). To avoid skewing of radiance data, graphical representation for each group was stopped after death of the first animal in the group.

### Statistical analysis

All comparisons between two groups were performed using either a two-tailed unpaired Student’s t-test or Mann-Whitney test, depending on normality of distribution. Comparisons between more than two groups were performed by two-way analysis of variance (ANOVA) with Bonferroni correction for multiple comparisons. All results are represented as mean ± standard error of the mean (s.e.m.). Survival data were analyzed using the Log-Rank (Mantel-Cox) test.

