## Supplementary figures and images for "Rational protein engineering to enhance MHC-independent T cell receptors"

### Supplementary Figure 1

#### Supplementary figure 1

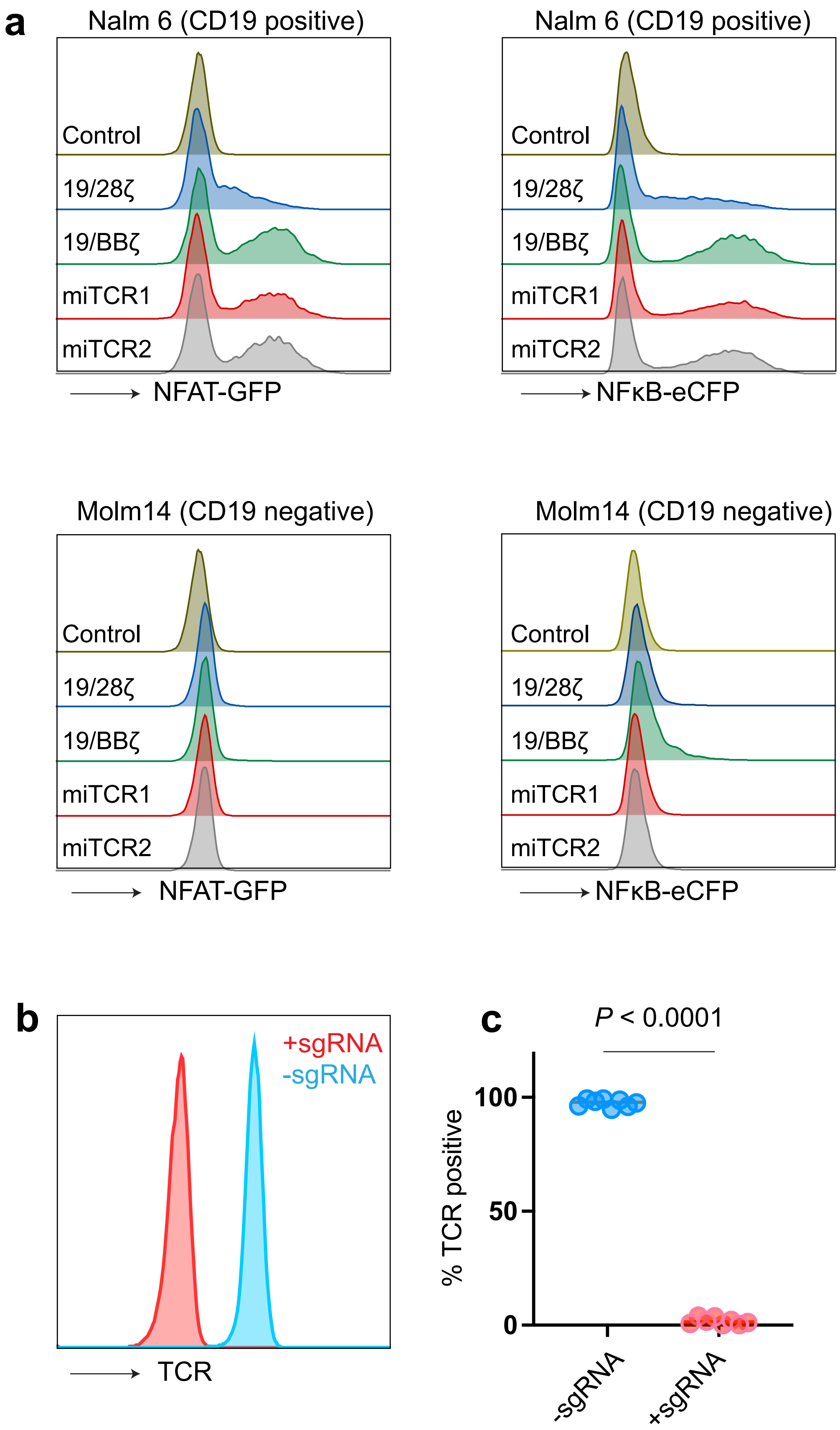

### Supplementary Figure 2

#### Supplementary figure 2

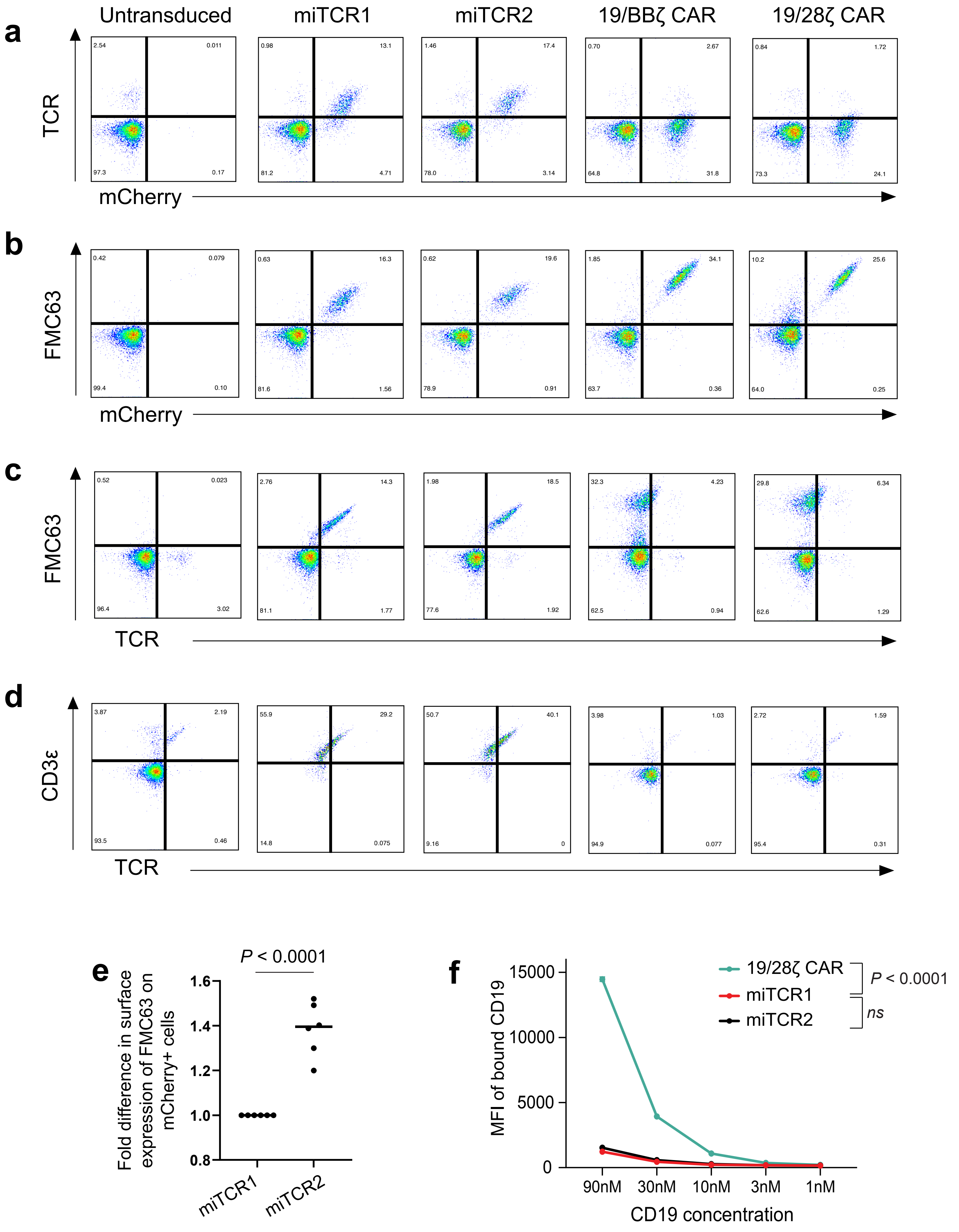

### Supplementary Figure 3

#### Supplementary figure 3

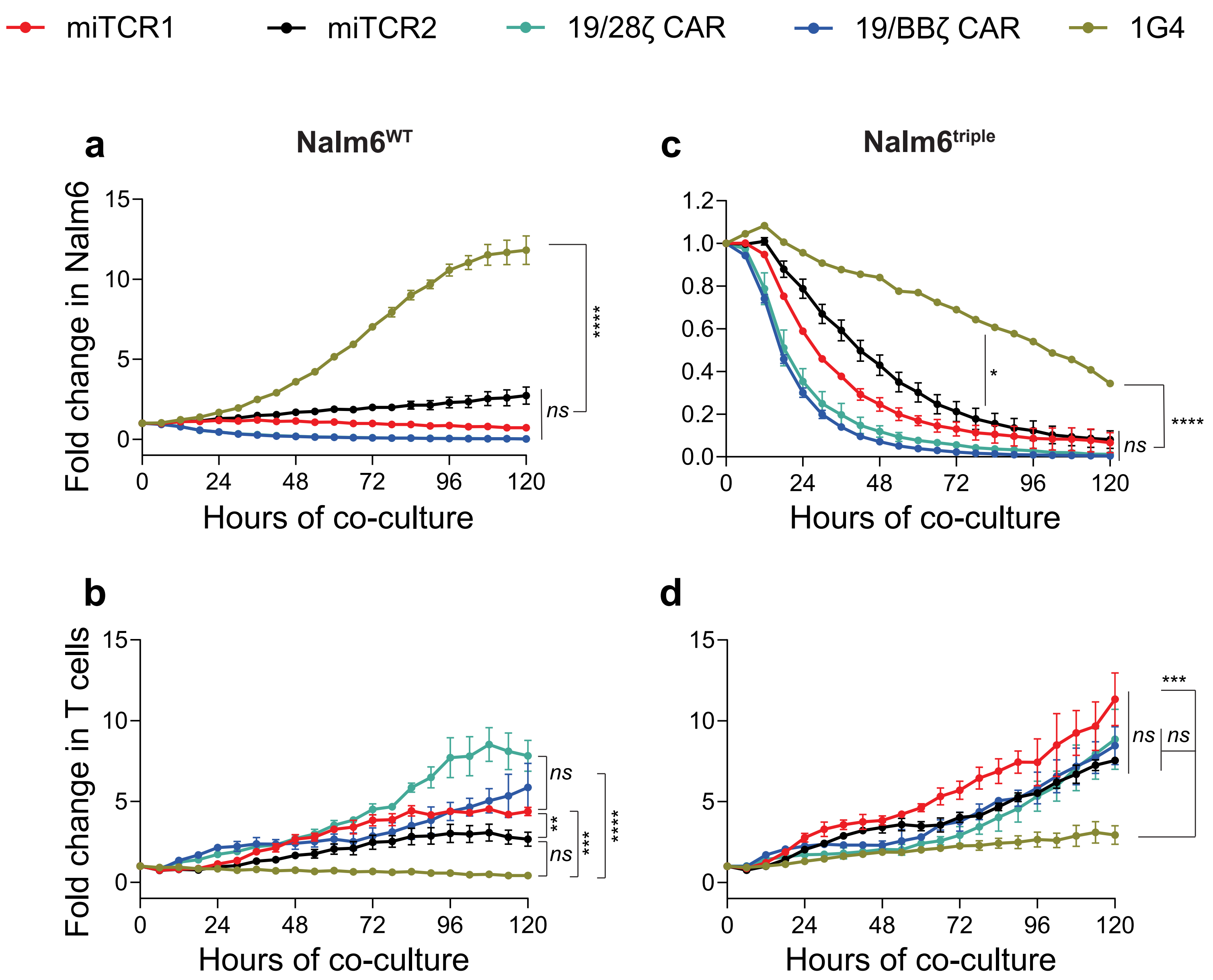

### Supplementary Figure 4

#### Supplementary figure 4

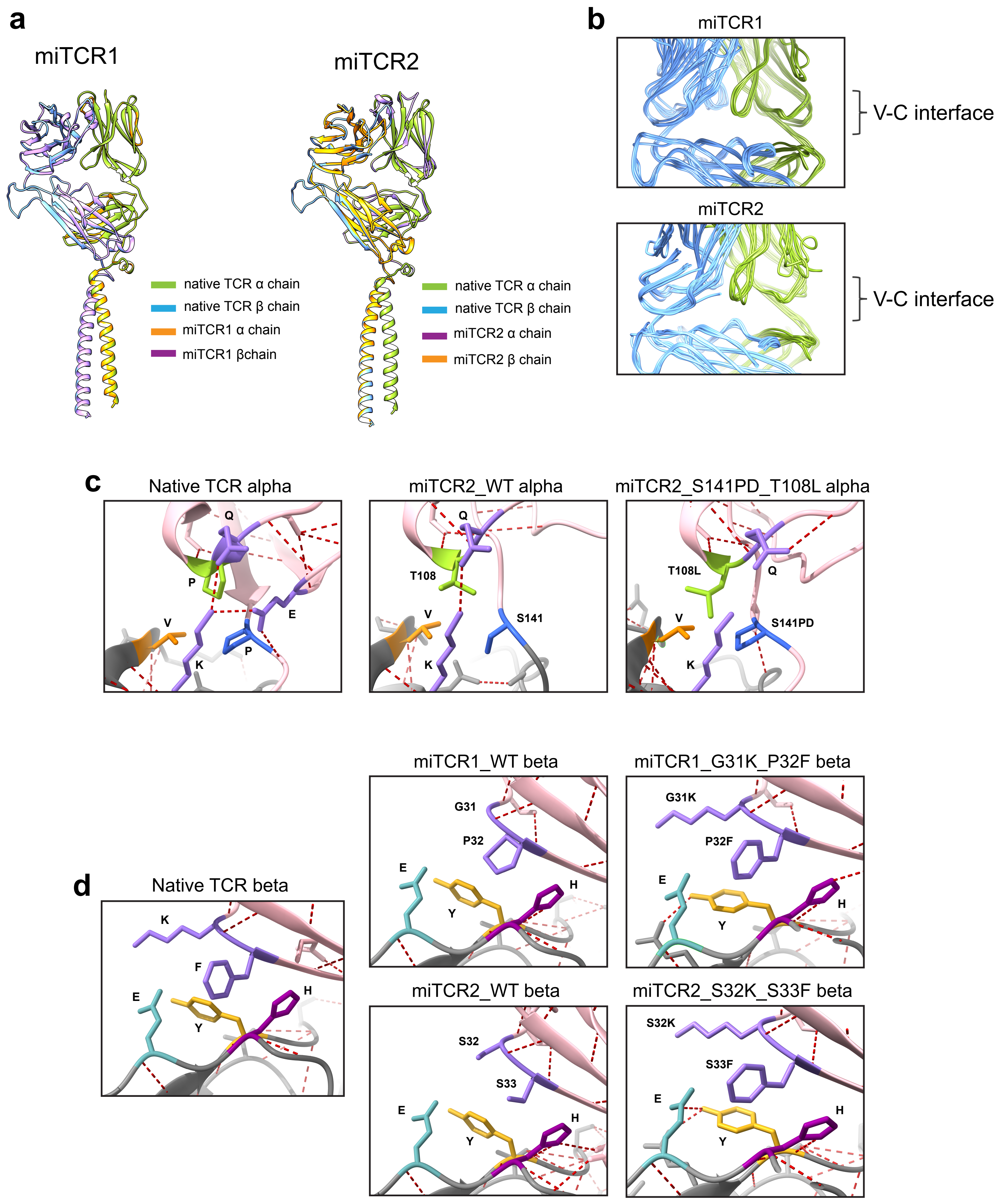

### Supplementary Figure 6

#### Supplementary figure 6

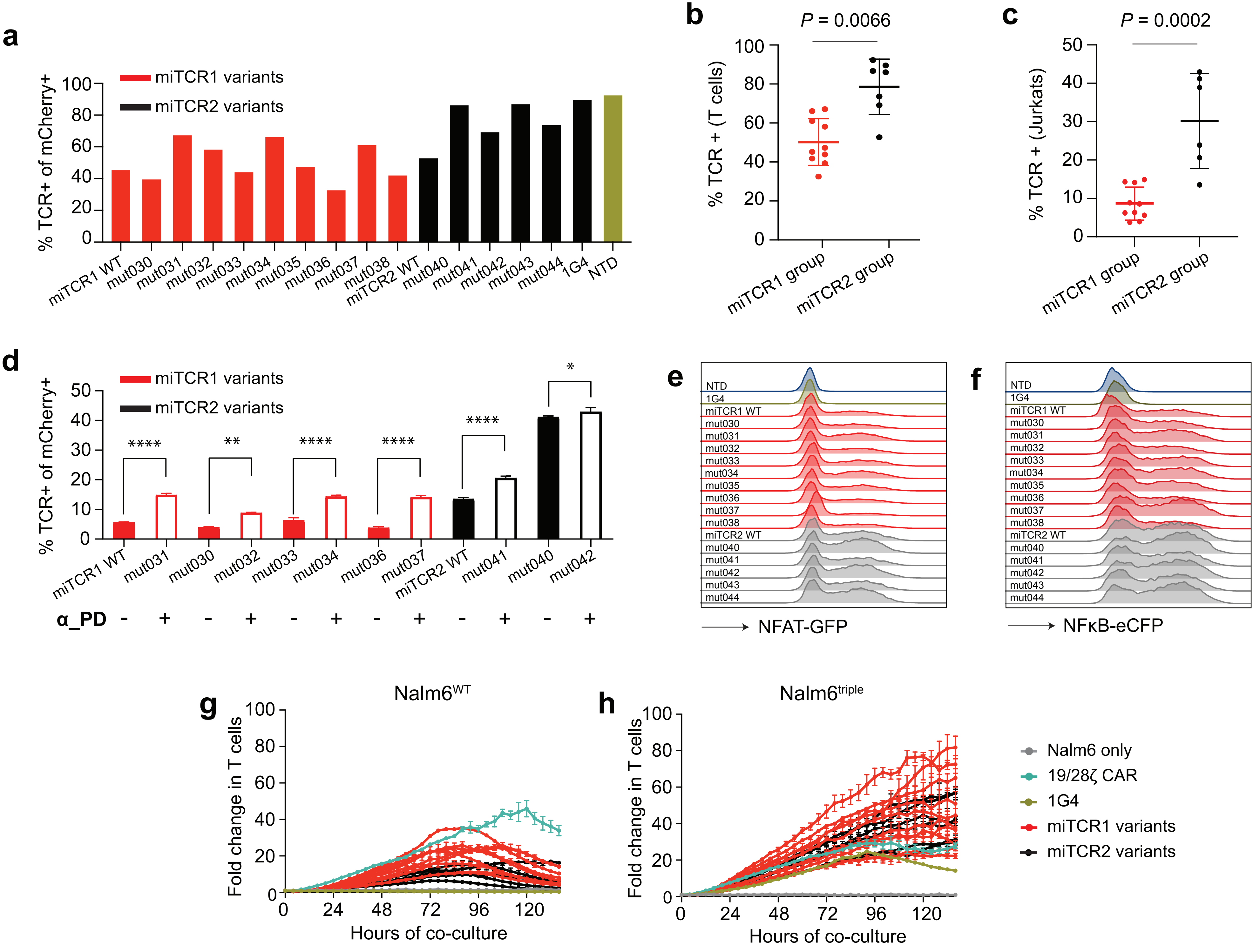

### Supplementary Figure 7

#### Supplementary figure 7

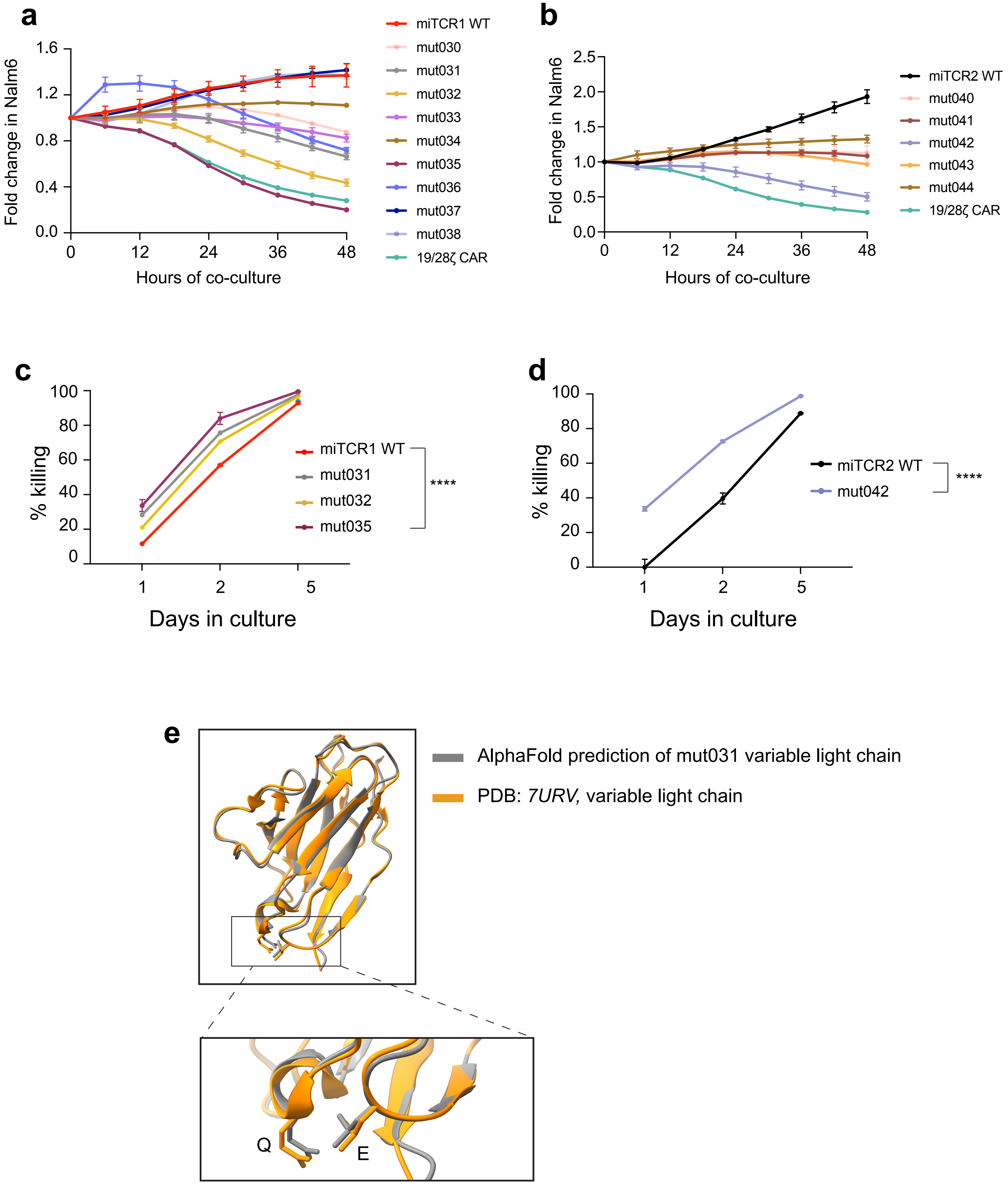

### Supplementary Figure 8

#### Supplementary figure 8

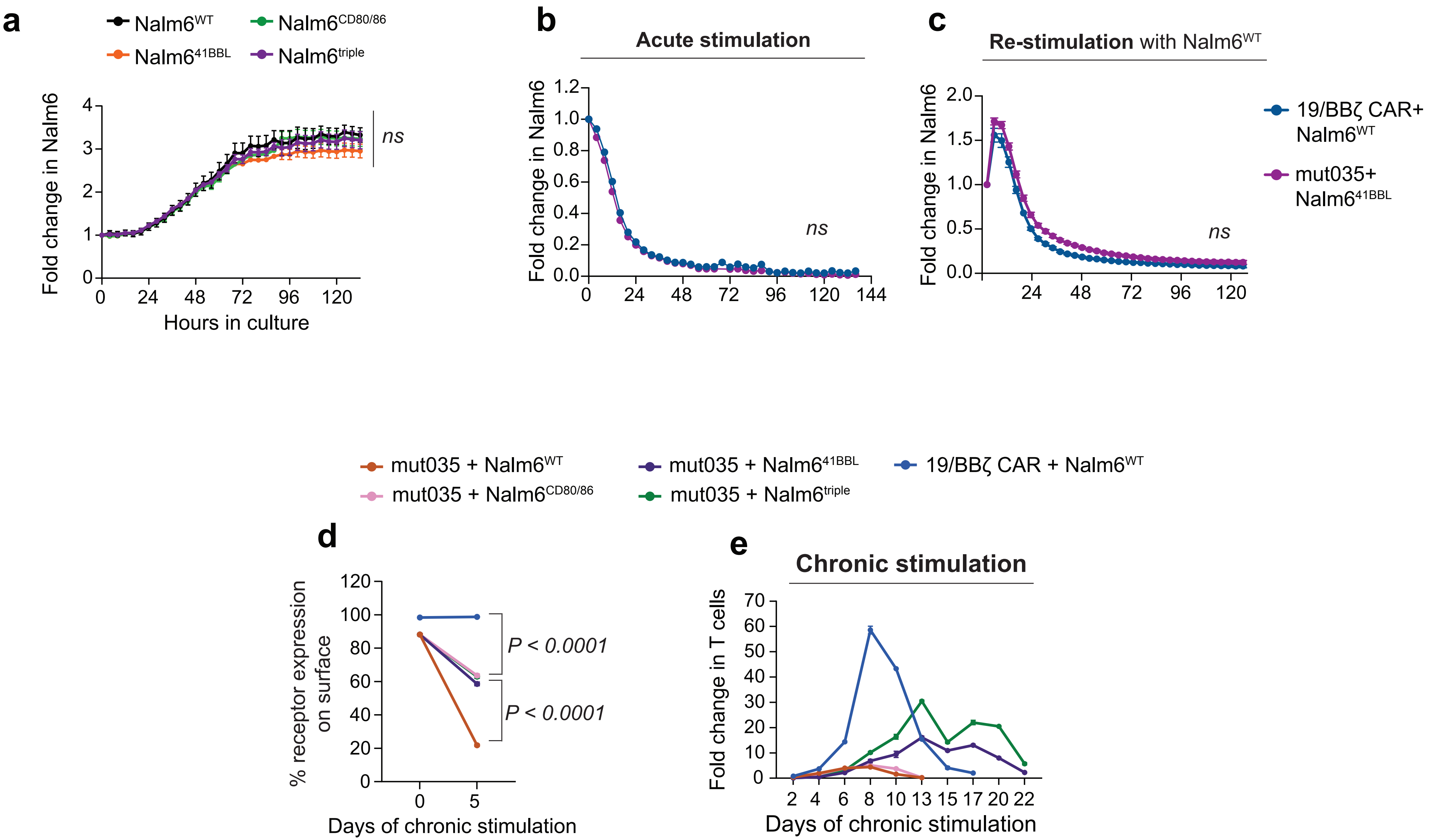
