## Supplementary Figure 5 for "Rational protein engineering to enhance MHC-independent T cell receptors"

| Name of receptor | Composition of receptor |
|---|---|
| miTCR1 WT | miTCR1_α_wt_β_wt |
| mut030 | miTCR1_α_wt_β_G31K_P32F |
| mut031 | miTCR1_α_PD_β_wt |
| mut032 | miTCR1_α_PD_β_G31K_P32F |
| mut033 | miTCR1_α_Q101L_β_wt |
| mut034 | miTCR1_α_PD_Q101L_β_wt |
| mut035 | miTCR1_α_PD_Q101L_L36E_β_wt |
| mut036 | miTCR1_α_Q101L_β_G31K_P32F |
| mut037 | miTCR1_α_PD_Q101L_β_G31K_P32F |
| mut038 | miTCR1_α_PD_Q101L_L36E_β_G31K_P32F |
| miTCR2 WT | miTCR2_α_wt_β_wt |
| mut040 | miTCR2_α_wt_β_S32K_S33F |
| mut041 | miTCR2_α_S141PD_β_wt |
| mut042 | miTCR2_α_S141PD_β_S32K_S33F |
| mut043 | miTCR2_α_S141PD_T108L_β_wt |
| mut044 | miTCR2_α_S141PD_T108L_β_S32K_S33F |
